# Design of a *Streptococcus pyogenes* M protein immunogen to elicit M type cross-reactivity

**DOI:** 10.1101/2021.12.07.471543

**Authors:** Kuei-Chen Wang, Eziz Kuliyev, Victor Nizet, Partho Ghosh

## Abstract

Coiled coil-forming M proteins of the widespread and potentially deadly bacterial pathogen *Streptococcus pyogenes* (Strep A) are immunodominant targets of opsonizing antibodies. However, antigenic sequence variability into >220 M types, as defined by the M protein hypervariable region (HVR), has been considered to limit its utility as a vaccine immunogen due to type-specificity in the antibody response. Surprisingly, a multi-HVR immunogen in clinical vaccine trials elicited M type cross-reactivity. The basis for this cross-reactivity is unknown but may be due in part to antibody recognition of a three-dimensional (3D) pattern conserved in many M protein HVRs that confers binding to human C4b-binding protein (C4BP). To test this hypothesis, we asked whether a single M protein immunogen carrying the 3D pattern would elicit cross-reactivity against other M types carrying the 3D pattern. We found that a 34-amino acid sequence of M2 protein bearing the 3D pattern retained full C4BP-binding capacity when fused to a coiled coil-stabilizing sequence from GCN4. This immunogen, called M2G, elicited cross-reactive antibodies against a number of M types that carry the 3D pattern but not against those that lack the 3D pattern. The M2G antiserum recognized M proteins as displayed natively on the Strep A surface, and promoted the opsonophagocytic killing of Strep A strains expressing these M proteins. As C4BP-binding is a conserved virulence trait of Strep A, targeting the 3D pattern may prove advantageous in vaccine design.

## Introduction

*Streptococcus pyogenes* (Group A *Streptococcus* or Strep A) is a globally widespread gram-positive bacterial pathogen that causes a variety of diseases, ranging from mild and self-limiting (e.g., pharyngitis and impetigo) to invasive and deadly (e.g., necrotizing fasciitis and streptococcal toxic shock syndrome) (1). Strep A infection can also lead to autoimmune diseases (e.g., acute rheumatic fever and rheumatic heart disease), which remain serious causes of morbidity and mortality in the developing world (2–4). Approximately 500,000 deaths occur annually due to diseases caused by Strep A (5). At present there is no vaccine against Strep A (6), with one of the major impediments being the sequence variability of its immunodominant surface antigen, the bacterial cell wall-anchored M protein (7–11).

More than 220 M protein types have been identified (12). The primary sequence of M proteins in general have heptad repeats, which are diagnostic of α-helical coiled coils (13), and structural studies have directly confirmed that M proteins do indeed form parallel, dimeric α-helical coiled coils (14–16), albeit with functionally significant sequence and structural irregularities (17, 18). The sequence of the N-terminal 50 amino acids of the mature form of M proteins (with their signal sequences removed) is hypervariable and defines the M type. These N-terminal hypervariable regions (HVRs) elicit protective, opsonizing antibodies (7–11). In contrast, other portions of M proteins are often not immunogenic or do not elicit opsonizing antibodies (7, 19). In addition, M protein HVRs do not elicit autoreactive antibodies (20), which other portions of M proteins do, presenting a concern for initiating autoimmune diseases (21). While M protein HVRs have highly favorable features as vaccine immunogens, antibody reactivity tends to be type-specific and therefore limited to a single M type strain (22–25).

Surprisingly, a Strep A vaccine immunogen composed of multiple M protein HVRs elicited an M type cross-reactive response. This vaccine immunogen, StreptAnova™, consists of 30 different M protein HVRs fused into four separate polyproteins (~45-50 kDa per polyprotein), and upon immunization of rabbits elicited reactivity against these 30 M types as well as cross-reactivity against ~50 M types not included in the immunogen (20, 26, 27). This reactivity promoted opsonophagocytic killing of Strep A (26, 27). The human immune system also appears to be capable of generating an M type cross-reactive response (28, 29). The basis for M type cross-reactivity of StreptAnova™ is not known. However, our own work in understanding how human C4b-binding protein (C4BP) binds multiple M protein HVRs provides a plausible model (16, 30).

C4BP limits the generation of the major opsonin C3b and thereby functions as a downregulator of the complement system (classical and lectin pathways). Recruitment of C4BP by M protein to the Strep A surface is an essential virulence trait, preventing opsonization, phagocytic uptake, and consequent killing (16, 31–38). A large-scale study found that 90 of 100 Strep A strains of differing M types bound C4BP (33). Because C4BP-binding has been attributed only to M protein HVRs (33), these results suggested that C4BP is cross-reactive for M protein HVRs. To understand the basis of M type cross-reactivity of C4BP, we determined X-ray crystal structures of four M protein HVRs (M2, M22, M28, and M49) each bound to C4BPα1-2, a fragment of C4BP that is necessary and sufficient to bind M protein HVRs (16). These structures revealed these M protein HVRs display a similar spatial or three-dimensional (3D) pattern of amino acids that contact a common site in C4BP. The amino acids of this shared 3D pattern are surrounded in space and in primary sequence by a larger number of variable amino acids, and so in effect the 3D pattern is diluted within the variability of the HVR (30). However, once the 3D pattern was identified, it was recognizable in the primary sequence of M proteins of about 40 of the ~90 Strep A strains (16) that were shown to bind C4BP (33).

Based on this observation, we hypothesized that the typical antibody response was M type-specific simply because variable amino acids outnumber those in the conserved 3D pattern. However, if an antibody were to bind amino acids of the conserved 3D pattern in one M type, it should then also recognize other M types that have this 3D pattern. Therefore, such an antibody would be M type cross-reactive. Notably, 15 of the 30 M protein HVRs in StreptAnova™ have the C4BP-binding 3D pattern, and correspondingly 20 of the ~50 cross-reactive M protein types elicited by StreptAnova™ have the 3D pattern (26, 27), suggesting that at least some of the cross-reactivity of StreptAnova™ is due to recognition of the 3D pattern. Likewise, M type crossreactivity observed for three other multi-HVR immunogens may be explained by recognition of the 3D pattern (39–41). The composition of these three immunogens, which are pentavalent or hexavalent and mostly contain HVRs that are also in StreptAnova™, is based on physicochemical properties rather than M type prevalence in North America and Europe as it is for StreptAnova™) (26). Together these results suggest that the C4BP-binding 3D pattern is capable of eliciting M type cross-reactive antibodies.

To test this hypothesis directly, we pursued a detailed proof-of-principle study. We used a short (34-amino acid) sequence from a single C4BP-binding M protein for immunization. M2 protein was chosen since its binding to C4BP has been studied in detail through mutagenesis (16). A 34-amino acid (aa) portion of M2 centered on the 3D pattern maintained full C4BP-binding affinity when fused to the canonical coiled-coil forming protein GCN4 (42). The antiserum evoked by the resulting immunogen, called M2G, was reactive against M2 and crossreactive against a number of C4BP-binding M types but not against M types that do not bind C4BP. The M2G antiserum was not cross-reactive against C4b or self-antigens. Reactivity and cross-reactivity of the M2G antiserum extended to M proteins displayed natively on the Strep A surface and resulted in the opsonophagocytic killing of Strep A strains.

## Results

### Minimized C4BP-binding regions of M2 protein

Our previous structural studies used M2^N100^ (Fig. 1a), a protein fragment consisting of the N-terminal 100 aa of the mature form of the protein (i.e., with its signal sequence cleaved), for co-crystallization with C4BPα1-2 (16). The structure revealed that the C4BP-binding region of M2 protein localized to a span of only 23 amino acids (aa 61-83) within the HVR (Fig. 1a). To limit immunoreactivity to the C4BP-binding amino acids of the M2 HVR, we first asked whether a short fragment of M2 protein constituting just the C4BP-binding amino acids would maintain C4BP binding. However, expression of M2 aa 61-83 by recombinant means in *E. coli* was poor and yielded insufficient quantities of protein for further experiments. We tried longer M2 fragments, either aa 42-86 (M2_42_) or 53-86 (M2_53_) (Fig. 1a). Amino acid 42 is the very N-terminus of mature M2 protein, and 53 and 86 are the first and last amino acids, respectively, that are ordered in the crystal structure of M2 bound to C4BPα1-2 (16). Both M2_42_ and M2_53_ were expressed recombinantly in sufficient quantities for further studies. However, neither M2_42_ nor M2_53_ bound His-tagged C4BPα1-2 above background levels (Figs. 1b, and S1a, b). A fragment of M22 protein (M22_248_, aa 42-248) was used as a positive control for C4BP-binding in this and other experiments. It seemed possible that M2_42_ and M2_53_ were too short to form a dimeric, α-helical coiled coil efficiently, a necessity for M protein to bind C4BP (16, 43). To overcome this problem, we fused short sequences from the ideal coiled-coil forming protein GCN4 (42) to M2 protein fragments, maintaining a continuous heptad register between the two (Table S1). We first tried sandwiching M2 aa 61-83 between single GCN4 heptads (Fig. 1a, GM2_61_G), but observed no binding to C4BP (Fig. S1c). Next, we tried longer GCN4 coiled-coil sequences of about three or four heptads (23 or 27 aa) fused to the C-terminus of M2 aa 53-86 or 61-83; these fusion constructs were called M2_53_G and M2_61_G, respectively (Table S1). While M2_61_G bound C4BPα1-2 slightly above background level, M2_53_G bound C4BPα1-2 well, with an affinity apparently higher than that of M22_248_ (Figs. 1b, c and S2a, b). For simplicity, we refer to M2_53_G as M2G hereafter.

**Figure 1.**
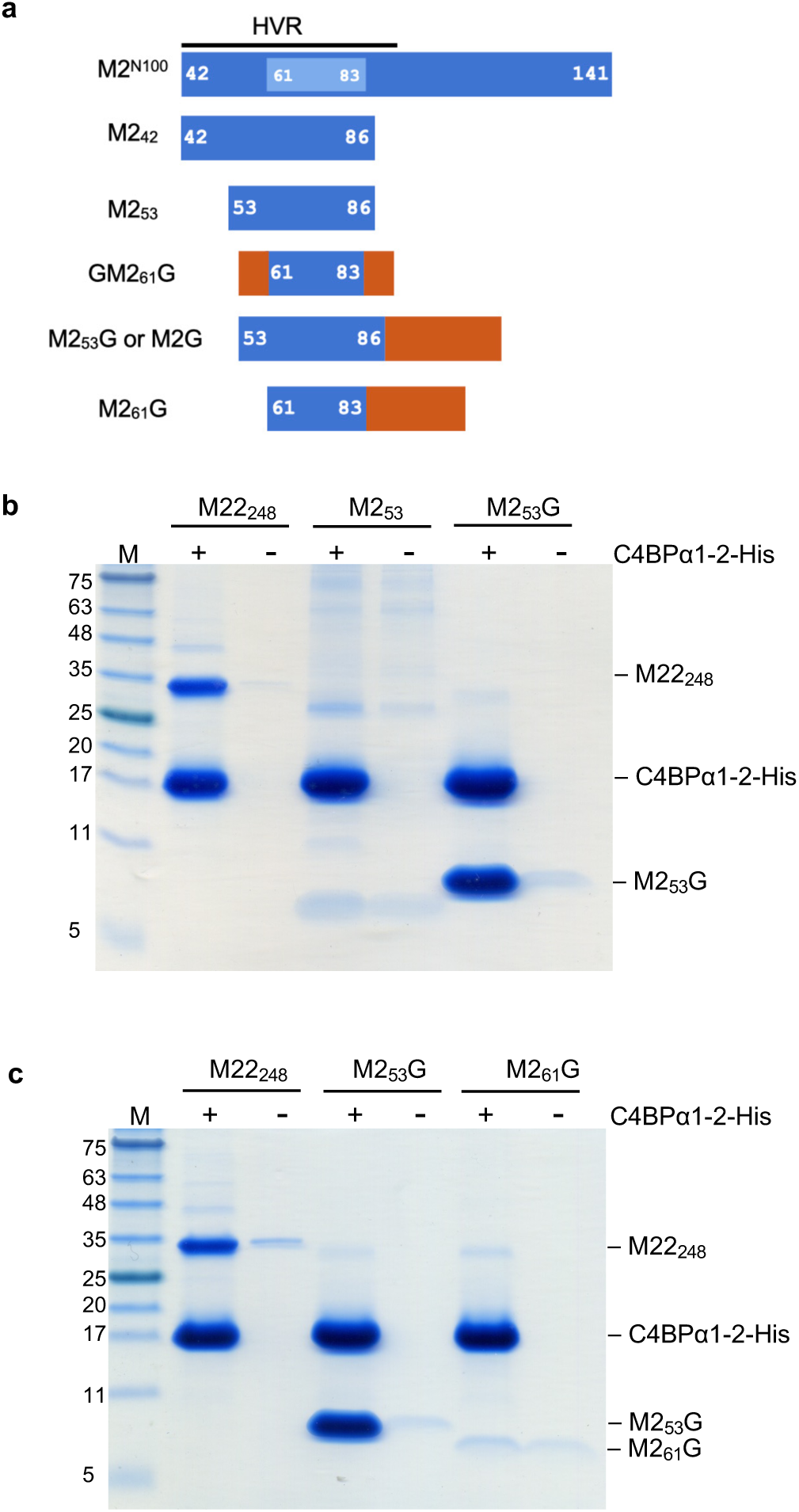
Binding of minimized M2 protein to C4BPα1-2. **a.** Schematic of M2 constructs. M2 in blue (with C4BP-contacting aa in cyan) and GCN4 in rust. **b and c.** Interaction of M2 protein constructs (b, M2_53_ and M2_53_G; c, M2_53_G and M2_61_G; M22_248_ was used as a positive control) with C4BPα1-2-His at 37 °C, as assessed by a Ni^2+^-NTA agarose co-precipitation assay and visualized by non-reducing, Coomassie-stained SDS-PAGE. Bound fractions are shown. Input samples are shown in Fig. S2. Each gel is representative of at least three experimental replicates.

We then asked whether M2G recapitulated the C4BP-binding affinity of intact M2 protein. Isothermal titration calorimetry (ITC) was carried out and showed that the K_D_ of C4BPα1-2 bound to intact M2 protein was identical to that of C4BPα1-2 bound to M2G, 4 μM (Fig. 2, Table 1). Thus, M2G, which contains only 34 aa of M2, possessed the full C4BP-binding affinity of intact M2 protein. Furthermore, these results suggested that GCN4 aided the coiled-coil dimerization of this M2 region to restore its C4BP binding ability.

**Figure 2.**
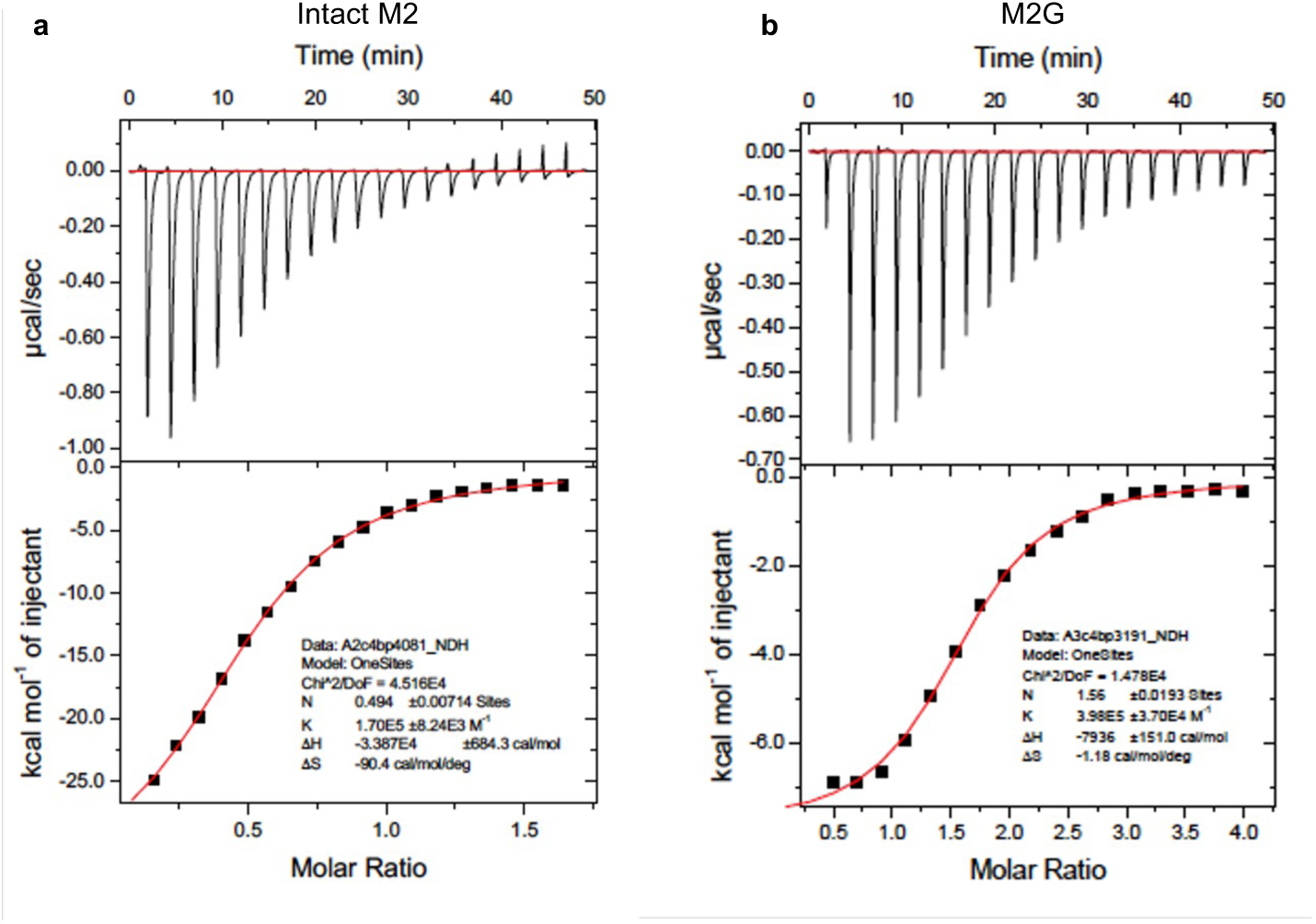
ITC Isotherms and Isograms. **a,** Intact M2 protein and **b,** M2G were titrated into a solution of C4BPα1-2. The top half of each panel shows isotherms, and the bottom half isograms. The binding curves were fitted using a single site model with the Origin software package. Each panel is representative of three experimental replicates.

**Table 1.** Isothermal titration calorimetry analysis of M protein-C4BPα1-2 interaction.

| | $K_D$ (μM) | $N^a$ | Stoichiometry<br>(C4BP : M) | $\Delta H$ (kcal/mol) | $-T\Delta S$ (kcal/mol) |
| --- | --- | --- | --- | --- | --- |
| **intact M2** | $4 \pm 2$ | $0.50 \pm 0.01$ | 2:1<sup>b</sup> | $-24 \pm 10$ | $17 \pm 10$ |
| **M2G** | $4 \pm 1$ | $0.51 \pm 0.01$ | 2:1 | $-27 \pm 6$ | $20 \pm 6$ |

<sup>a</sup> $N$ , binding stoichiometry of the M protein.
<sup>b</sup> Two molecules of C4BPα1-2 per one M protein dimer.

### M2G as an immunogen

Having identified that M2G recapitulated C4BP-binding, we asked whether it was sufficient to evoke an immune response that was cross-reactive against M protein types that carry the 3D pattern. Rabbit polyclonal antibodies were raised against M2G and assayed for reactivity against various recombinant M proteins. While M2, M22, M28 and M49 HVRs all present similar 3D patterns of amino acids that are complementary to C4BP, these spatial patterns are exhibited differently in the heptad repeats of their primary sequences (16). The heptad patterns of M2 and M49 HVRs are similar to one another and belong to one subset, the M2/M49 sequence pattern; and the M22 and M28 HVR patterns are similar to one another and belong to a second subset, the M22/M28 sequence pattern (16). We chose M protein types from each pattern that are prevalent in human infectious disease epidemiology (44, 45). For the M2/M49 group, these were M2, M49, M73, M77, and M89 proteins, and for the M22/M28 group, these were M4, M11, M22, M28, M44, and M81 proteins. As negative controls, we used M1, M5, and M6 proteins, which do not bind C4BP and lack the 3D pattern.

We expressed and purified constructs constituting the N-terminal 100 amino acids of the mature forms of these M proteins. Binding to C4BP had not been directly evaluated for some of the M proteins. For these an enzyme-linked immunosorbent assay (ELISA) for C4BP-binding was carried out (Fig. S3). All of the M^N100^ constructs (i.e., N-terminal 100 aa of the mature form) belonging to the M2/M49 or M22/M28 pattern bound C4BP, except for M77^N100^, which bound C4BP at the level of the negative control M5^N100^ construct.

We next tested the reactivity and cross-reactivity of the M2G antiserum. As expected, the M2G antiserum recognized M2^N100^ well, with an antibody titer that was significantly greater than that of pre-immune serum (>10^5^ vs. <10^2^) (Figs. 3 and S4). The M2G antiserum was cross-reactive with titers > 10^3^ against all the M^N100^ constructs belonging to the M2/M49 pattern (i.e., M49, M73, and M89), except for M77^N100^, which as noted above did not bind C4BP. For the M22/M28 pattern, the M2G antiserum was cross-reactive with a titer > 10^3^ against only M28^N100^. Statistically significant cross-reactivity was also seen for M11^N100^, but the titer was low (< 10^3^). For the remaining members of M22/M28 group (M22^N100^, M4^N100^, M44^N100^, and M81^N100^), the titer of the M2G antiserum was low (< 10^3^) and not significantly different from that of the pre-immune serum (Fig. 3 and S4). The titer of the M2G antiserum was uniformly low against M^N100^ constructs of M proteins that are known not to bind C4BP (i.e., < 10^3^ for M1, and < 10^2^ for M5 and M6) (Figs. 3 and S4). These results indicate that a single M protein antigen can elicit crossreactivity based on the C4BP-binding 3D pattern.

**Figure 3.**
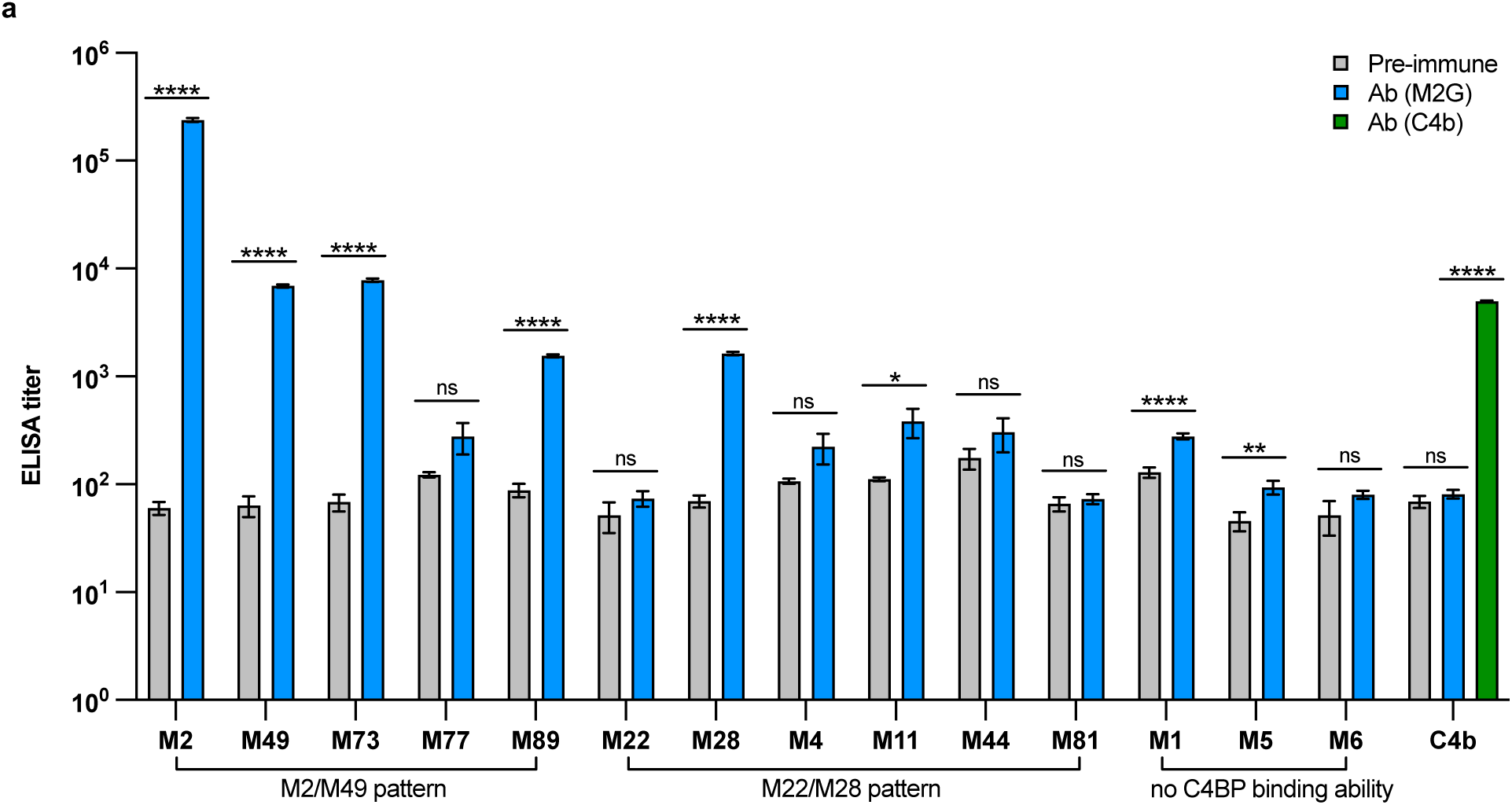
Reactivity and cross-reactivity of M2G antiserum. Titers of pre-immune and immune M2G sera against M^N100^ constructs (N-terminal 100 amino acids of the mature form of the protein) and C4b, as determined by ELISA. M^N100^ constructs and C4b were adhered to ELISA plate wells. Mean and standard errors of the mean are shown. All experiments were carried out in triplicate and performed at least two times. Statistical analyses were performed by Student’s *t*-tests and one-way ANOVA for M^N100^ constructs and C4b, respectively; *p* < 0.05 *, *p* < 0.01 **, *p* < 0.001 ***, *p* < 0.0001 ****, *p* > 0.05 (not significant, ns).

Because both M protein HVRs and C4b are bound by nearly the same site on C4BP (31), we asked whether the M2G antiserum possessed unwanted cross-reactivity against C4b. The M2G antiserum did not cross-react against C4b, as evaluated by ELISA with C4b adhered to a solid substrate (Fig. 3, titer < 10^2^). The conformational integrity of C4b adhered to the solid substrate was verified by its recognition by an anti-C4b antibody (Figs. 3 and S4c). These results are consistent with the observation that M protein HVRs and C4b differ in their binding mode for C4BP (31).

While autoreactivity is not attributed to M protein HVRs, this remains a general concern for vaccines based on M proteins (46). To evaluate the reactivity of the M2G antiserum against human tissues affected in Strep A autoimmune sequelae, western blot analysis was performed with normal adult human brain tissue lysate (HB) and heart tissue lysate (HH). Because autoreactivity can be due to portions of M proteins outside of the HVR, we also raised rabbit antibodies against intact M2 protein and compared the cross-reactivity of the M2G antiserum against that of the intact M2 protein antiserum. The M2G antiserum reacted against intact M2 protein but not HB or HH (Fig. S5a). In contrast, the antiserum raised against intact M2 protein reacted against intact M2 and both HB and HH (Fig. S5b). These results suggest that the M2G immunogen does not elicit cross-reactivity against human tissues, whereas intact M2 protein has the potential to do so. These results are also consistent with M protein HVRs not eliciting autoreactive antibodies (20).

We asked whether the pattern of M protein cross-reactivity described above was limited to a single rabbit or reproducible in a second rabbit. To this end, a second rabbit was immunized with M2G and the cross-reactivity of this second rabbit’s M2G antiserum was examined (Figs. S6 and S7). The reactivity and cross-reactivity patterns were almost identical with a Pearson correlation coefficient of 0.998. Some small differences were evident. For example, for the M2/M49 pattern, the relative cross-reactivity to M89^N100^ was lower in the second rabbit than in the first, and in the M22/M28 pattern, low-titer cross-reactivity to M44^N100^ but not M11^N100^ was statistically significant. Experiments were continued using the antiserum from the first rabbit.

As the M2G antigen contains sequences from both M2 protein and GCN4, we asked whether cross-reactivity was due to antibodies specific for the M2 as opposed to the GCN4 portion. We focused on M protein constructs that yielded the highest titers (> 10^3^), namely M2^N100^, M49^N100^, M73^N100^, M89 ^N100^, and M28^N100^ (Fig. 3). Reactivity of the M2G antiserum to these M protein constructs was competed with increasing concentrations of either M2G or M6G (Table S1). The latter consists of a portion of M6 protein (aa 56-89) fused to nearly the identical portion of GNC4 in M2G (Table S1). For all M^N100^ constructs except for M49^N100^, competition occurred with M2G but not M6G, even when M6G was used at >2-fold higher concentration (Fig. S8a). The integrity of M6G as a competitor was demonstrated by adhering His_6_-tagged M6G to ELISA plates and using soluble His_6_-tagged M6G as a competitor (Fig. S8b). These results indicated that for M2^N100^, M73 ^N100^, M28 ^N100^, and M89 ^N100^, cross-reactive antibodies were specific to the M2 portion of the M2G immunogen.

For M49^N100^, both M2G and M6G competed against the M2G antiserum for binding (Fig. S8a), suggesting that some or all of the cross-reactive antibodies against M49^N100^ were specific to the GCN4 portion of the M2G. M49 protein was eliminated from further studies.

### Strep A surface binding and opsonophagocytic activity of M2G antiserum

To assess whether the M2G antiserum recognized M proteins on the bacterial surface, we carried out flow cytometry on whole, living Strep A strains of differing M types. In line with results using purified proteins, the M2G antiserum bound the surface of an M2 strain to a significantly higher extent than did the pre-immune serum (Fig. 4, Table S2). Likewise, the M2G antiserum displayed significant cross-reactivity compared to the pre-immune serum against Strep A M73, M89, and M28 strains. No cross-reactivity was seen against an M5 strain, which does not bind C4BP. Overall, these results confirm that the M2G antiserum recognizes M proteins in their native conformation on the Strep A surface.

**Figure 4.**
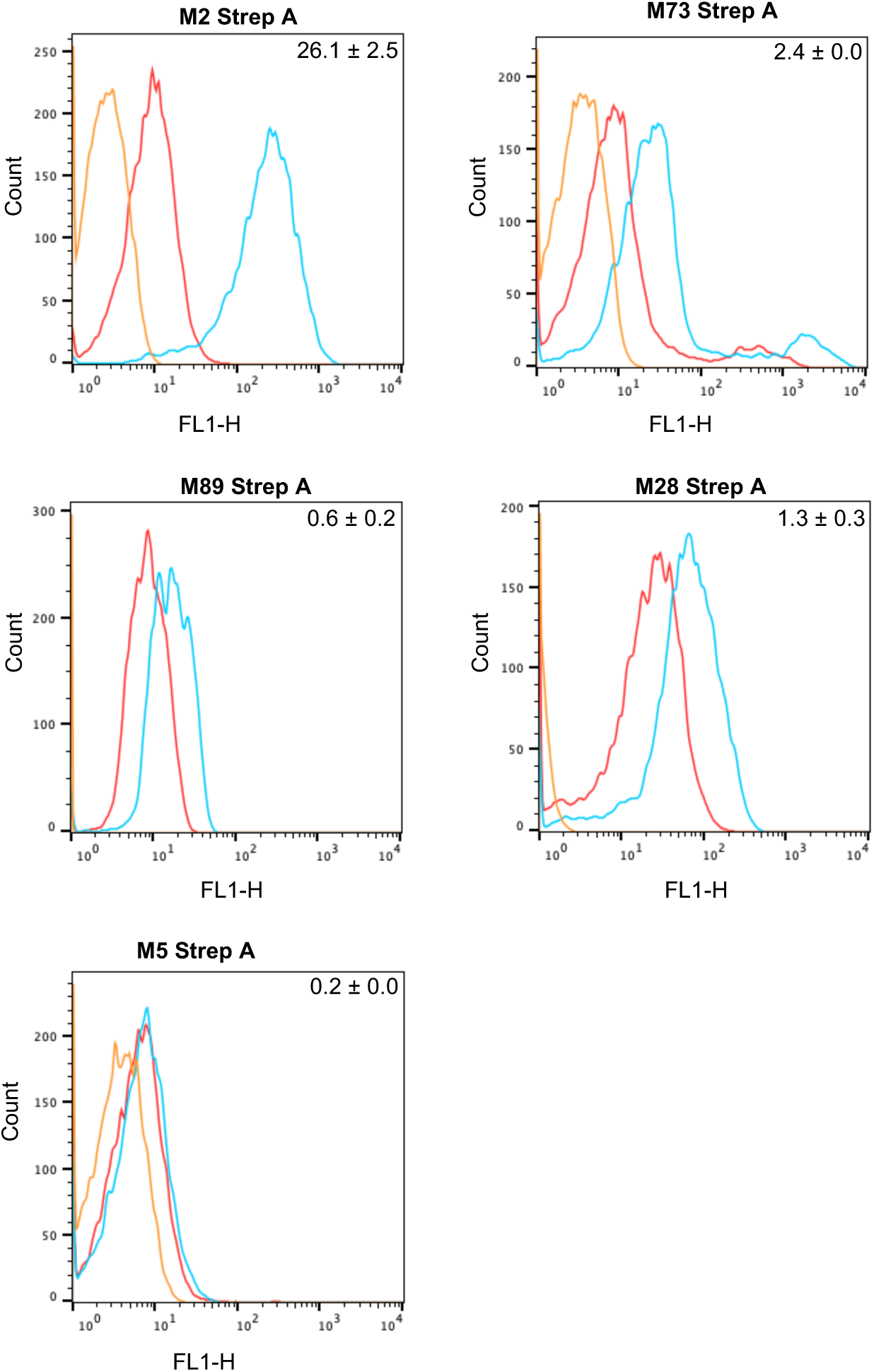
M2G antiserum binding to Strep A. Binding of pre-immune serum and M2G antiserum to Strep A M2, M73, M89, M28, and M5 strains, assessed by flow cytometry. Histograms show fluorescent intensities from pre-immune serum (red), M2G antiserum (blue), and anti-rabbit IgG antibody control (orange, 2° antibody only). Each histogram is representative of three experimental replicates. Numbers on the top right corner of each panel is the difference between the geometric mean of the fluorescent signal of the M2G antiserum and that of pre-immune serum, normalized by that of the pre-immune serum. The fluorescent signal of pre-immune serum (pre) and M2G antiserum (Ab(M2G)) are listed in Table S2.

Antibodies against M protein HVRs elicit opsonophagocytic antibodies (7–11). To verify that this is the case for the M2G antiserum, we evaluated whether the M2G antiserum promoted opsonophagocytic killing (OPK) of reactive and cross-reactive Strep A strains. For the OPK assay, we used cultured HL-60 cells differentiated to have a neutrophil-like phenotype, along with baby rabbit serum as the source of complement (47). While neutrophil-like HL-60 cells are not nearly as potent killers as primary neutrophils (48), they offer several advantages over the classical Lancefield assay (24). First and foremost, individual variation in complement and neutrophil activity is eliminated in the HL-60 assay, as is the existence of immunity against various M types (49). The HL-60 assay has been proposed as a standard means for evaluating Strep A vaccine candidates (47, 49). We focused on the M types against which the greatest reactivity and cross-reactivity had been demonstrated — M2, M73, M89, and M28 — and used an M5 strain as a negative control. As a positive control for the assay, we used IVIG, a concentrated pool of human antibodies. We found that IVIG was most potent against M28 and M73 strains (100% and 94.6 ± 0.2% killing) followed by M2 (72.1 ± 4.5%), M89 (52.7 ± 4.2%), and M5 (30.6 ± 3.2%) strains (Fig. 5).

**Figure 5.**
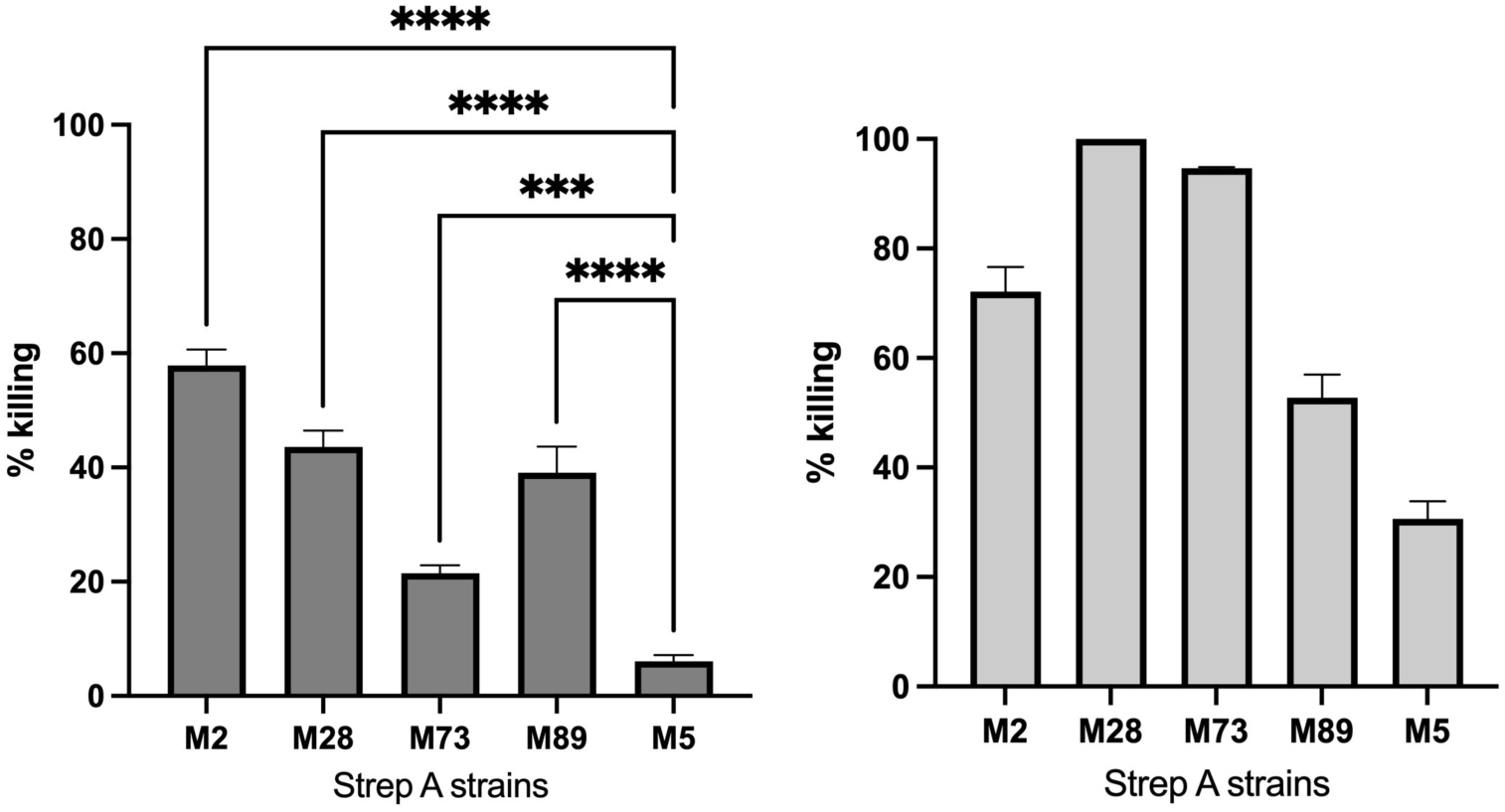
Bactericidal activity of M2G antiserum. Opsonophagocytic killing promoted by M2G antiserum (left) or IVIG (right) of Strep A M2, M28, M73, M89 and M5 strains. The assay was performed with DMF-differentiated HL-60 cells and baby rabbit complement. All experiments were carried out in triplicate and performed three times. Statistical analysis for the M2G antiserum was performed by one-way ANOVA; *p* < 0.05 *, *p* < 0.01 **, *p* < 0.001 ***, *p* < 0.0001 ****, *p* > 0.05 (not significant, ns).

Substantial OPK activity was found for the M2G antiserum. The highest OPK activity was against the M2 strain (57.9 ± 2.8 %) followed by the M28 (43.6 ± 2.9%), M89 (39.1 ± 4.5%), and M73 strains (21.5 ± 1.4%). These values were statistically significant compared to the OPK activity against the M5 strain (6.1 ± 1.1%). These results confirmed that antibodies directed against M protein HVRs promote opsonophagocytic killing.

## Discussion

We set out to test the hypothesis that directing the antibody response to the C4BP-binding 3D pattern in one M protein type would lead to cross-reactivity against other M protein types that share the 3D pattern. The design was to have the 3D pattern constitute as great a proportion of the immunogen as possible, and correspondingly, variable amino acids as minimal as possible. The well-studied M2 protein was chosen for these studies (16). Short segments of M2 protein containing the 3D pattern (e.g., aa 53-86) bound C4BPα1-2 poorly, likely because they formed unstable coiled coils. This finding may reflect the presence of charged amino acids at some of the core *a* positions of the coiled-coil heptad (16). Stabilization of coiled-coil structure through fusion of these M2 segments to portions of GCN4 (i.e., in M2G) restored C4BP binding. While the 3D pattern in M2 protein appeared to start at aa 61 (16), inclusion of amino acids upstream of this position had a significant effect on C4BP binding, leading to a K_D_ that matched that of intact M2 protein. In the crystal structure, these upstream amino acids contact a crystallographically related C4BPα1-2 molecule (16). Based on these results, it is likely that the contacts made by these upstream amino acids to C4BP are not a crystallization artifact but instead a *bona fide* interaction.

The M2G immunogen elicited a cross-reactive response against M73 and M89 proteins from the M2/M49 pattern. Cross-reactivity was also observed against M49 protein, but this was in part or entirely attributable to antibodies that were specific to GCN4 rather than M2 portions in the M2G immunogen. No notable sequence similarity exists between M49 and GCN4, and thus the basis for this surprising result requires further investigation. No cross-reactivity was seen to M77 protein, which despite having an unambiguous M2/M49 sequence pattern (16), did not bind C4BP. In the case of some M proteins (50), C4BP binding is significantly enhanced by the concurrent binding of Fc domains from human IgG. M77 protein may require this additional interaction to bind C4BP. Alternatively, C4BP binding may be conferred by a portion of M77 protein outside the HVR or another bacterial surface-associated protein. The M2G immunogen elicited a strong cross-reactive against only one member of the M22/M28 pattern, M28 protein. While the spatial arrangement of C4BP-interacting amino acids (i.e., 3D pattern) is similar for both M2/M49 and M22/28 sequence patterns (16), the different positioning of M protein α-helices between these two patterns appears to influence cross-reactivity. Clearly, further work is required to increase the scope of cross-reactivity of a 3D pattern-based immunogen. It may be worthwhile in the future to explore the use of consensus C4BP-binding sequences from the M2/M49 and M22/M28 patterns rather than the sequence of a single M type, and a means for stabilizing the coiled coil without fusion to GCN4 or other proteins.

The M2G antiserum cross-reacted against M proteins in their native conformation on the Strep A surface, which was likely favored through the use of an immunogen that retained C4BP-binding and hence native conformations. Furthermore, the M2G antiserum promoted the opsonophagocytic killing of Strep A strains of multiple M types, consistent with results showing that M protein HVRs evoke opsonic antibodies (7–11). The range of killing of Strep A by HL-60 cells due to cross-reactivity seen here (22-44%) was similar or slightly better than that observed in another study (16-41%), in which peptides derived from multiple M protein HVRs served as the immunogen (51). Notably, the cross-reactivity we observed correlated better with C4BP-binding than sequence identity (Table S3). For example, M77 protein, which did not bind C4BP and was not recognized by the M2G antiserum, has 59% identity with the M2G immunogen, but M28 protein, which did bind C4BP and was recognized, has only 35% identity. Opsonophagocytic killing of the M28 Strep A strain at 44% was the highest of all the cross-reactive interactions observed.

These results provide evidence that the conserved C4BP-binding 3D pattern can elicit antibodies that cross-react against M protein types that have the 3D pattern, and promote the opsonophagocytic killing of such Strep A strains. Significantly, the recruitment of C4BP to the Strep A surface is an essential virulence trait for numerous Strep A strains (35, 36, 38), and thus, escape from a broadly protective antibody that targets the C4BP-binding 3D pattern through further sequence variation may be limited by pressure to maintain C4BP interaction during infection (30). In effect, the C4BP-binding 3D pattern is an Achilles’ heel of many M protein types. These results provide impetus to pursue further experiments aimed at optimizing an immunogen based on the C4BP-binding 3D pattern.

## Materials and Methods

### Streptococcus pyogenes

The following *S. pyogenes* strains, which are clinical isolates from the U.S. Centers for Disease Control and Prevention, were used in this study: *emm2* (strain 3752-05), *emm5* (strain 3292-05), *emm28* (strain 4039-05), *emm49* (strain 3487-05), *emm73* (strain 3962-05) and *emm89* (strain 4264-05). *S. pyogenes* was grown statically in Todd-Hewitt broth (THB, BD) supplemented with 1% yeast extract (Gibco) overnight at 37 °C, and afterwards subcultured in the same medium until mid-logarithmic growth phase (OD_600_ = 0.4-0.6).

### Cloning and DNA manipulation

Coding DNA sequences for intact mature M1, M2, M4, M5, M22, M28 and M49 proteins were cloned, as described previously (14, 16), from *S. pyogenes* strains M1 (strain 5448), M2 (AP2), M4 (Arp4), M5 (Manfredo), M22 (Sir22), M28 (strain 4039-05), and M49 (NZ131), respectively, and ligated into pET-28b vector (Novagen) or a modified pET-28a vector (Novagen) that had encoded an N-terminal His_6_-tag followed by a PreScission protease (GE Healthcare) cleavage site. Truncated forms of these proteins were subcloned from these vectors. The coding DNA sequence for GCN4 was subcloned from *Saccharomyces cerevisiae*. The coding sequence of the N-terminal 100 amino acids of M6, M73, M77, M89, M11, M44 and M81 proteins were chemically synthesized (Integrated DNA Technologies, Inc.) and inserted into the aforementioned modified pET-28a vector. M protein-GCN4 fusion constructs were produced by strand overlap extension PCR, and ligated into the modified pET-28a vector. Protein sequences of M protein-GCN4 fusion constructs are listed in Table S1.

### Protein expression and purification

M proteins were expressed in *Escherichia coli* BL21 (Gold) and purified as previously described (14, 16), except that imidazole was not included in the lysis buffer. C4BPα1-2 was expressed in *E. coli* Rosetta 2 (Novagen). The protein was purified and refolded as previously described (52) with minor modifications. Specifically, bacteria were lysed with a C-5 Emulsiflex (Avestin). After refolding and dialysis, C4BPα1-2 was applied to a HiTrap Q HP column (GE Healthcare) and eluted using a 0-1 M NaCl gradient in 50 mM Tris, pH 8.5.

### Co-precipitation assays

His_6_-tagged C4BPα1-2 (150 μg) was mixed with M protein constructs (molar ratio 1:1.2) in 50 μl phosphate-buffered saline (PBS) at 37 °C for 30 min under rotation. Ni^2+^-NTA agarose beads (100 μl of 50 % slurry), pre-equilibrated with PBS, were then added to the protein mix and incubated at 37 °C for 40 min under rotation. The beads were washed three times with 0.5 mL of PBS supplemented with 15 mM imidazole, and eluted with 40 μl of PBS supplemented with 300 mM imidazole. Proteins in the input and eluted fractions were resolved by non-reducing SDS-PAGE and visualized by Coomassie Staining.

### Isothermal titration calorimetry (ITC)

ITC experiments were performed at 23 °C on a ITC200 microcalorimeter (MicroCal, MA) with PBS as the assay buffer. Titrations were carried out with 300-500 μM intact M2 or M2G, which was loaded in the injection syringe (40 μl), and 30-50 μM C4BPα1-2, which was loaded in the sample cell (~250 μl). A typical titration experiment consisted of 19 injections of 2 μl over a duration of 4 s; each injection was separated by 150 s. The cell stirring speed was 1000 rev/min. Raw data were collected, and binding curves were fitted using a single site model with Origin software (MicroCal).

### Rabbit polyclonal antisera

Rabbit polyclonal antisera were raised commercially (Pocono Rabbit Farm & Laboratory (PRF&L), Canadensis, PA) against 200 μg of purified M protein constructs (M2G or intact M2 protein). An initial immunization in Complete Freund’s Adjuvant (CFA) was carried out, followed by 3 boosts with 100 μg purified protein in Incomplete Freund’s Adjuvant (IFA) on days 14 and 28, and 50 μg purified protein in IFA on day 56. A large bleed was performed on day 70 to obtain serum.

### ELISA

#### Determination of antibody titers

Purified M^N100^ protein constructs at 1 μg/mL were coated in the wells of 96-well microtiter plates (Corning) in carbonate buffer (50 mM Na_2_CO_3_-NaHCO_3_, pH 9.6) overnight at 4 °C. All subsequent procedures were performed at RT. Wells were washed three times in TBST (150 mM NaCl, 50 mM Tris, pH 8.0, and 0.1% Tween-20) and blotted dry after the preceding step and after all steps described below. Wells were blocked with 0.1% BSA in TBST for 1 h. They were then incubated with 100 μl of rabbit pre-immune or immune serum (serially diluted in 0.1% BSA/TBST) for 1.5 h. In the case of competition experiments, varying concentrations of M2G or M6G were incubated with the immune serum for 10 min prior to addition to wells. One hundred μl horseradish peroxidase (HRP)-conjugated goat anti-rabbit IgG (H+L) (Southern Biotech) at 1:4000 dilution in 0.1% BSA/TBST was added and incubated for 1 h. For detection, 100 μl TMB substrate (BD Biosciences) was added and incubated for 10 min (protected from light), followed by addition of 50 μl of 2 N sulfuric acid to stop the reaction. For determination of titers, the absorbance at 450 nm (A_450_) was measured and fit to a sigmoidal curve using GraphPad Prism. Antibody titers were defined as the reciprocal of the interpolated serum dilution level that yielded 50% of the maximum A_450_. Statistical analysis was performed using the Student’s *t*-test to compare immune and pre-immune sera.

#### Detection of C4b

An ELISA was carried out as above, except that C4b (Millipore) at 1 μg/mL was coated in ELISA plate wells, and detected by incubation for 1 h with 100 μl anti-C4b polyclonal antibodies (ThermoFisher) at 1:4000 dilution in 0.1% BSA/TBST, followed by incubation for 1 h with goat anti-chicken IgY-HRP (Santa Cruz Biotechnology) at 1:2500 dilution in 0.1% BSA/TBST.

#### His_6_-M6G vs His_6_-M6G competition

ELISAs were carried out as above, except for the following. His_6_-tagged M6G at 1 μg/mL in PBS was coated in the wells of 96-well microtiter plates. Soluble His_6_-tagged M6G at varying concentrations was incubated for 10 min with HRP-conjugated mouse anti-His antibody (BioLegend) diluted 1:2000 in 0.1% BSA/TBST. Wells were then incubated for 1.5 h with 100 μl of this solution.

#### M protein-C4BP interaction

ELISAs were carried out as above, except for the following. Intact C4BP (Complement Technology) at 10 μg/mL in PBS was coated in the wells of 96-well microtiter plates. Wells were incubated for 1.5 h with 10 μg/mL of His_6_-tagged M proteins (N-terminal 100 amino acids, diluted in 1% BSA/TBST). Wells were then incubated with 100 μl HRP-conjugated mouse anti-His antibody (1:2000 dilution in 1% BSA/TBST, BioLegend).

### Human tissue cross-reactivity

Twenty μg of normal adult human brain tissue lysate (Novus Biologicals) or heart tissue lysate (Novus Biologicals) was resolved on 4-20% gradient SDS-PAGE (Bio-Rad) and transferred to a PVDF membrane (Millipore) for immunoblotting. Membranes were blocked with 5% BSA in TBST at RT for 1 h, and then incubated with rabbit antisera (1:1000 dilution in 5% BSA/ TBST) at RT for 1 h. Membranes were washed three times by TBST for 5 min each. Membranes were then incubated with HRP-conjugated goat anti-rabbit IgG (H+L) (1:4000 dilution; Southern Biotech) at RT for 1 h, and SuperSignal west pico chemiluminescent substrate (Thermo Fisher Scientific) was then added. The resulting chemiluminescence was recorded on an ChemiDoc XRS+ imaging system (Bio-Rad).

### Antiserum binding to *S. pyogenes*

*S. pyogenes* was grown to mid-logarithmic phase, washed in PBS, and blocked with 10% heat-inactivated donkey serum in PBS (Sigma-Aldrich) at RT for 1 h. Heat-inactivated M2G antiserum or pre-immune serum was added to *S. pyogenes* to 1% final volume and incubated at RT for 1 h. After washing once in PBS, samples were incubated in 1:200 dilution of donkey antirabbit IgG antibody with Alexa Fluor 488 conjugation (BioLegend) at RT for 30 min (protected from light). Samples were then washed once in PBS, resuspended in PBS, and analyzed by flow cytometry (BD FACSCalibur). Fluorescent signal intensity was analyzed using FlowJo software (Tree Star Inc.).

### HL-60 opsonophagocytic killing assay

The OPK assay was performed as previously described (47, 49) with some modifications. HL-60 cells (CCL-240; ATCC) were cultured in RPMI medium (RPMI 1640 with 1% L-glutamine (Corning) and 10% heat-inactivated fetal bovine serum (Gibco)). Differentiation into neutrophil-like cells was carried out through incubation for four to five days and with a cell density of 4 x 10^5^ cells/mL in RPMI medium supplemented with 0.8% dimethylformamide (DMF). The phenotype of differentiated HL-60 cells was assessed by flow cytometry using mouse antihuman CD35 PE conjugated antibody (BioLegend) and mouse anti-human CD71 APC conjugated antibody (BioLegend). Differentiated cells were used in the OPK assay if >55% of cells were CD35^+^ and <15% of the cells were CD71^+^. Prior to use in the assay, differentiated HL-60 cells were washed first in Hank’s balanced salt solution (HBSS) without Ca^2+^/Mg^2+^ (Gibco) and then in HBSS with Ca^2+^/Mg^2+^ (Gibco), and resuspended at a concentration of 1 x 10^7^ cells/mL in fresh opsonization (OPS) buffer (HBSS with Ca^2+^/Mg^2+^, 0.1% gelatin, 5% heat-inactivated pig serum (Sigma-Aldrich), 1 mg/mL human fibrinogen (Millipore), and 10 U/mL heparin (Sigma-Aldrich)).

Prior to carrying out the OPK assay, *S. pyogenes* strains were passaged through HL-60 cells, as follows. *S. pyogenes* was grown to mid-logarithmic phase and then diluted in THB to 3,000-10,000 CFU/ml. Ten μl of *S. pyogenes* were incubated with 50 μL heat-inactivated normal rabbit serum (NRS, PRF&L) per well in a round-bottom 96-well plate (Corning) at RT for 30 min, followed by the addition of 40 μl of active baby rabbit complement (BRC, PelFreez) and 100 μl of differentiated HL-60 cells to each well. The plate was then sealed with aluminum film (AlumaSeal II AF100; Excel Scientific) and incubated at 37 °C for 2 h with end-over-end rotation. The final concentration of BRC in the reaction mixture was 5-20% in OPS buffer, with the specific value dependent on the *S. pyogenes* strain (such that non-specific killing, as described below, was <35%). After 2 h incubation, the plate was placed on ice for 30 min to stop the activity of HL-60 cells. After mixing thoroughly, 10 μl from each well was spotted on THB agar plates, which were tilted immediately to spread the bacteria in drips across the plates. The plates were incubated overnight at 37 °C, and *S. pyogenes* colonies were recovered. This procedure was carried out a second time to yield a total of two passages for each strain.

The OPK assay was carried out as above with twice-passaged *S. pyogenes*. Bacteria were incubated with HL-60 cells, heat-inactivated NRS, and heat-inactivated BRC (control A); HL-60 cells, heat-inactivated NRS, and active BRC (control B); or HL-60 cells, 50 μl heat-inactivated M2G antiserum or 50 μl heat-inactivated IVIG (Intravenous Immune Globulin (Human) 10%, Octapharma USA Inc.), and active BRC. After overnight incubation of THB agar plates, CFUs were enumerated.

The percentage of killing was calculated as ((CFU of control B – CFU of M2G antiserum or IVIG) / CFU of control B) x 100. The percentage of non-specific killing was calculated as (CFU of control A – CFU of control B) / CFU of control A) x 100. The number of bacterial generations was calculated by comparing the total CFU of control B to the CFU in the inoculum. Only assays in which the level of non-specific killing was <35%, the number of bacterial generations in control B was >4, and the CFU of controls A and B were between 50-200 were considered. Statistical analysis was performed using one-way ANOVA to compare M2, M28, M73 and M89 strains to M5.

## Acknowledgements

We thank Satoshi Uchiyama for assisting with flow cytometry analysis, and Nina J. Gao and Sanaz Salehi for advice on OPK assays. This work was supported by NIH R21AI140436 (P.G. and V.N.) and R01AI154149 (P.G.).

## Supplemental Figure Legends

**Figure S1.**
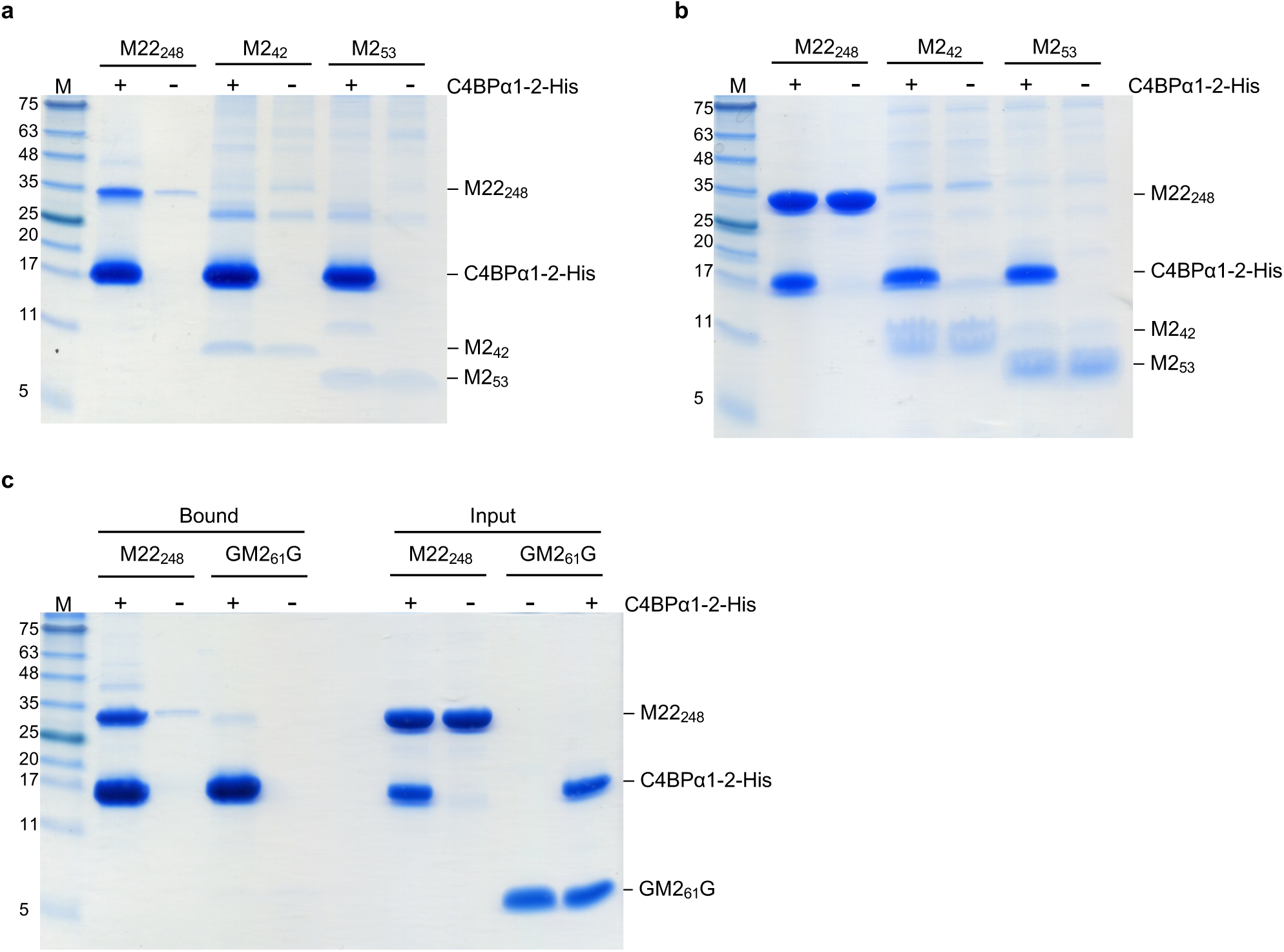
Binding of M2 protein constructs to C4BPα1-2. **a.** Interaction of M2_42_ and M2_53_ with C4BPα1-2-His at 37 °C, as assessed by a Ni^2+^-NTA agarose co-precipitation assay and visualized by non-reducing, Coomassie-stained SDS– PAGE. M22_248_ was used as a positive control. **b.** Input samples from panel a. **c.** Interaction of GM2_61_G with C4BPα1-2-His, carried out as in panel a. Each gel in the figure is representative of at least three experimental replicates.

**Figure S2.**
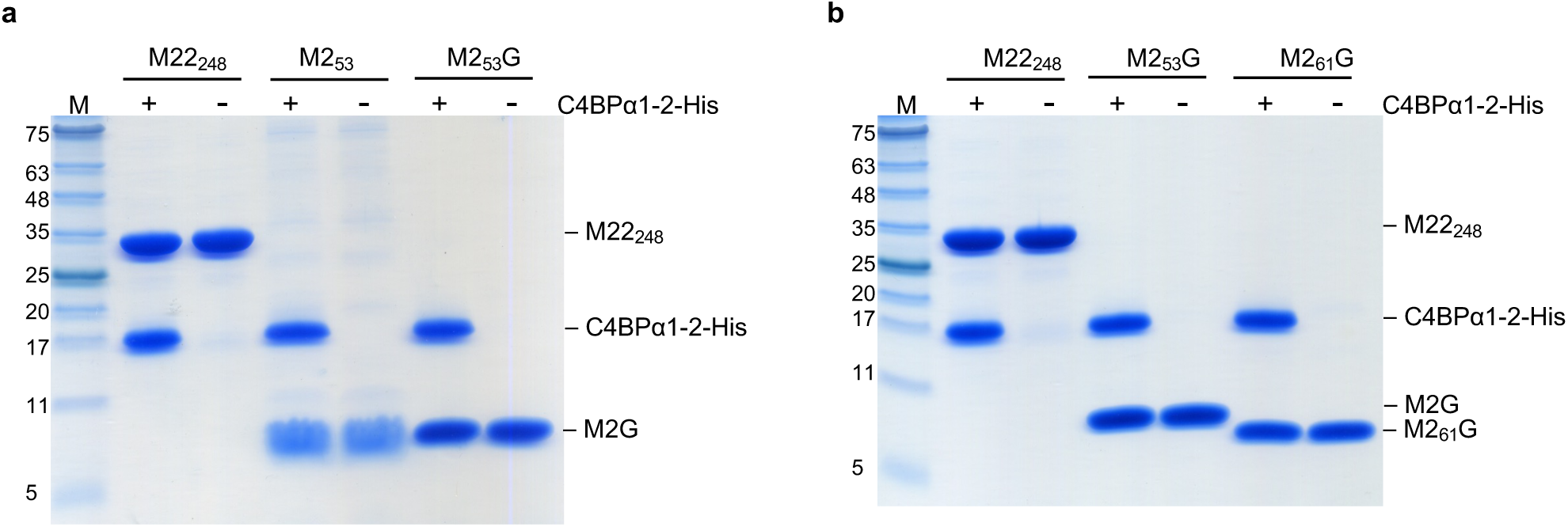
Binding of minimized M2 protein to C4BPα1-2. Input samples from Ni^2+^-NTA agarose co-precipitation experiments shown in Figures 1b and c visualized by non-reducing, Coomassie-stained SDS-PAGE (**a,** M2_53_ and M2_53_G; **b,** M2_53_G and M2_61_G; M22_248_ was used as a positive control in panels a-b). Each gel is representative of at least three experimental replicates.

**Figure S3.**
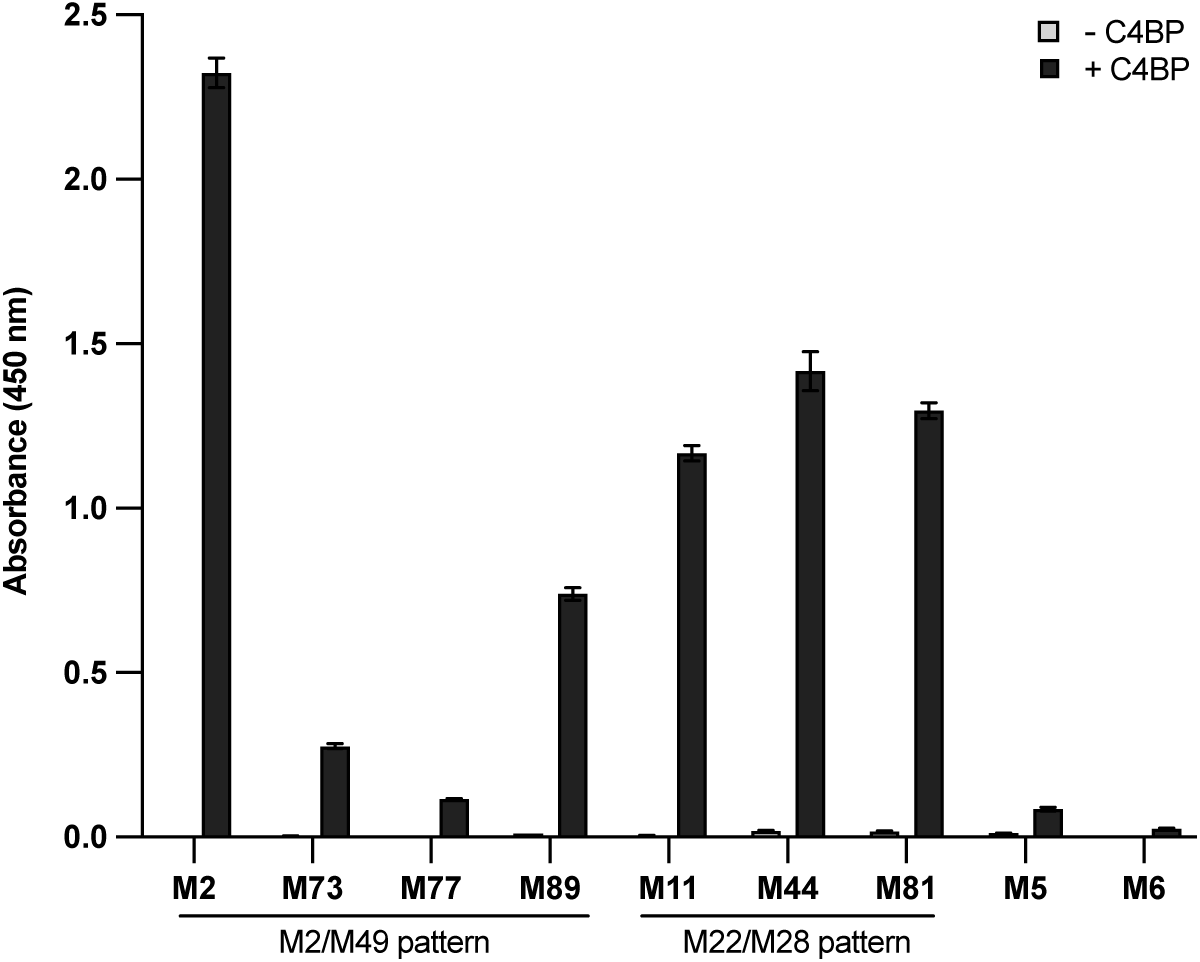
Binding of M proteins to C4BP. Interaction of His-tagged M^N100^ proteins (N-terminal 100 amino acids of mature form) with intact C4BP as assessed by ELISA. C4BP was adhered to the ELISA plate (+C4BP) or not adhered as a negative control (-C4BP), and M proteins were added and detected using an anti-His antibody. All experiments were carried out in triplicate and performed three times.

**Figure S4.**
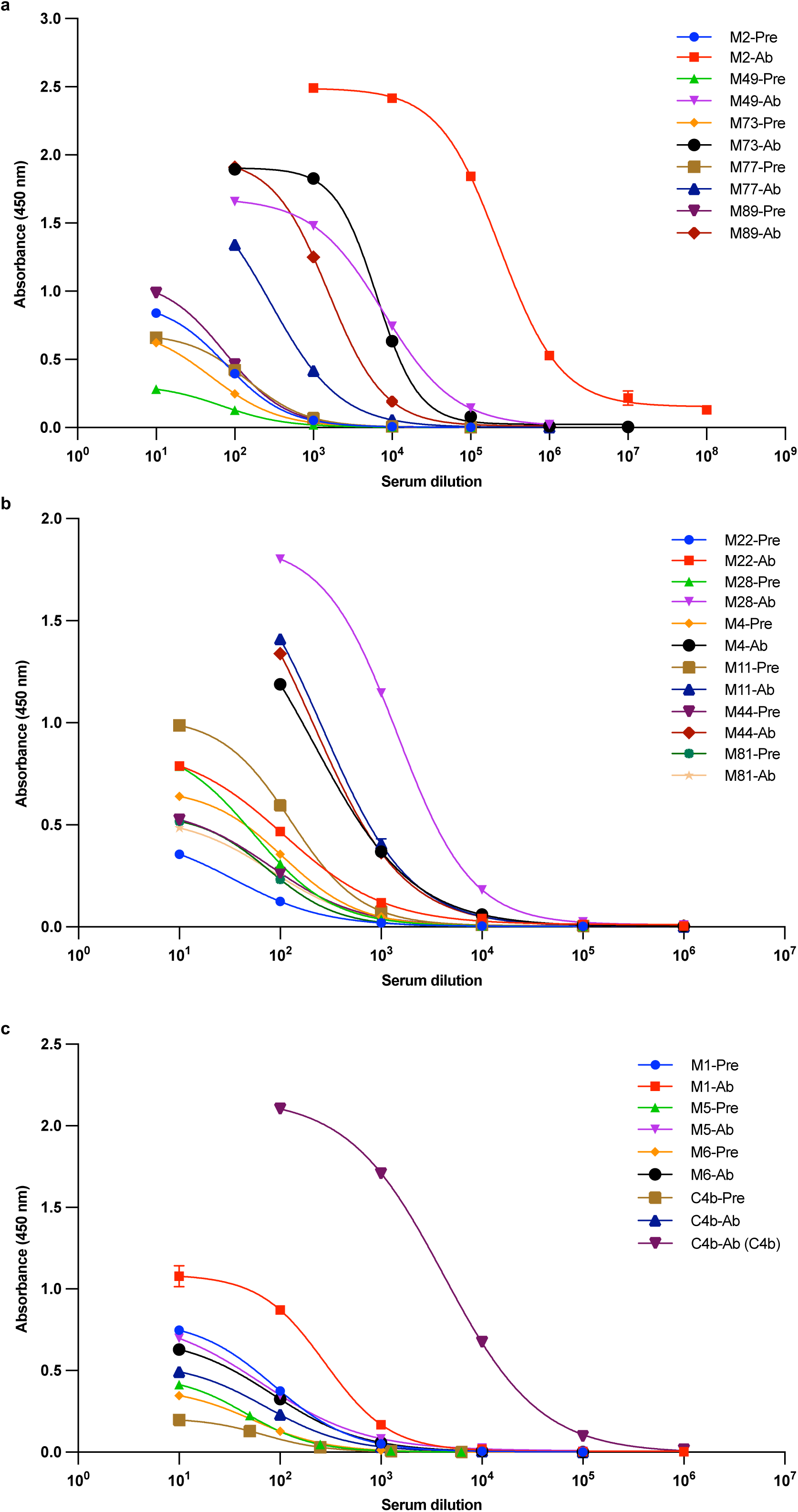
Titration curves of M2G antiserum against M^N100^ constructs and C4b. Titration curves fitted to binding of pre-immune serum (Pre) or M2G antiserum (Ab) to M^N100^ constructs (N-terminal 100 amino acids of the mature form of the protein) or C4b. **a,** M proteins belonging to the M2/M49 pattern. **b,** M proteins belonging to the M22/M28 pattern. **c,** C4b and M proteins that do not bind C4BP. Each curve is representative of at least two experimental replicates.

**Figure S5.**
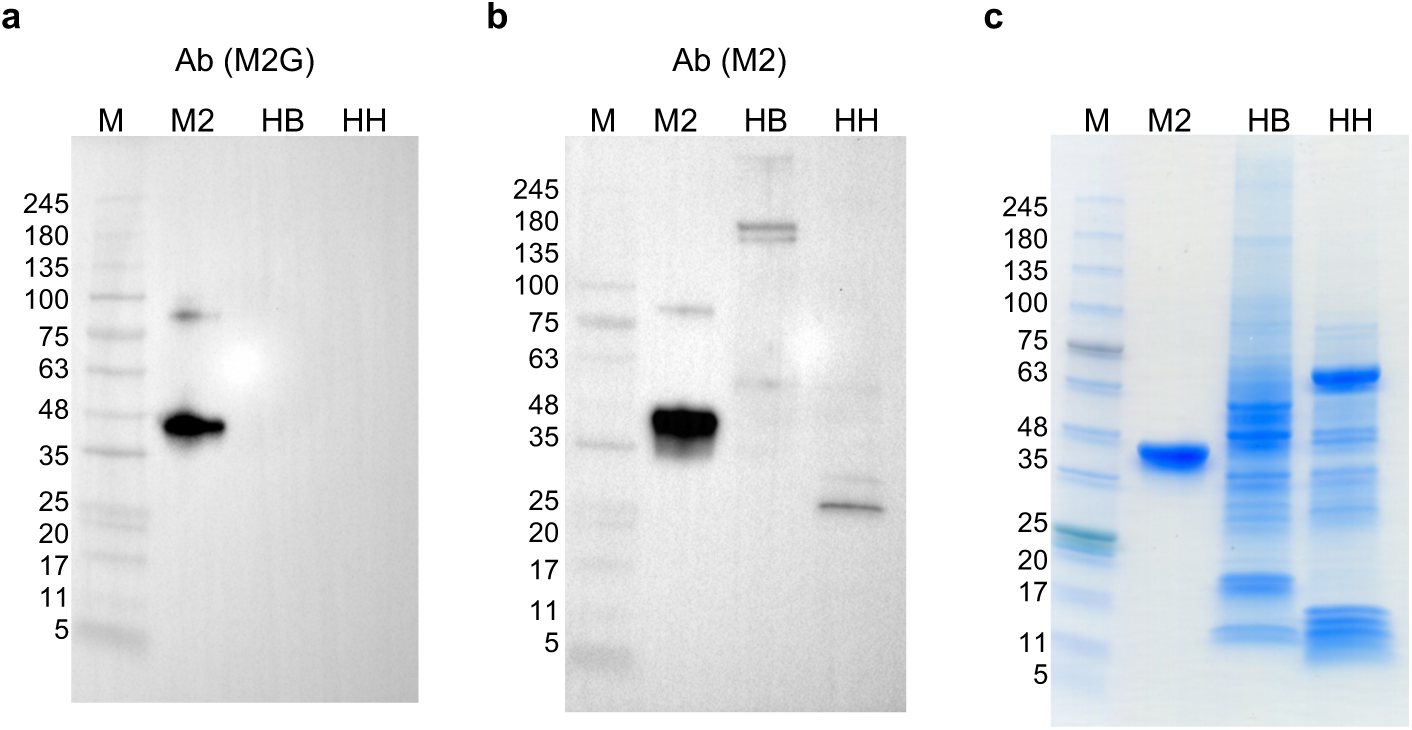
Reactivity of M2G antiserum against human tissues. Reactivity of M2G and M2 antisera against normal adult human brain tissue lysate (HB) or heart tissue lysate (HH), as determined by western blot analysis. Intact M2 protein (M2) was used as a positive control. Each blot is representative of three experimental replicates. **a,** M2G antiserum (Ab(M2G)). **b,** M2 antiserum (Ab(M2)). **c,** Input samples visualized by Coomassie-stained SDS-PAGE.

**Figure S6.**
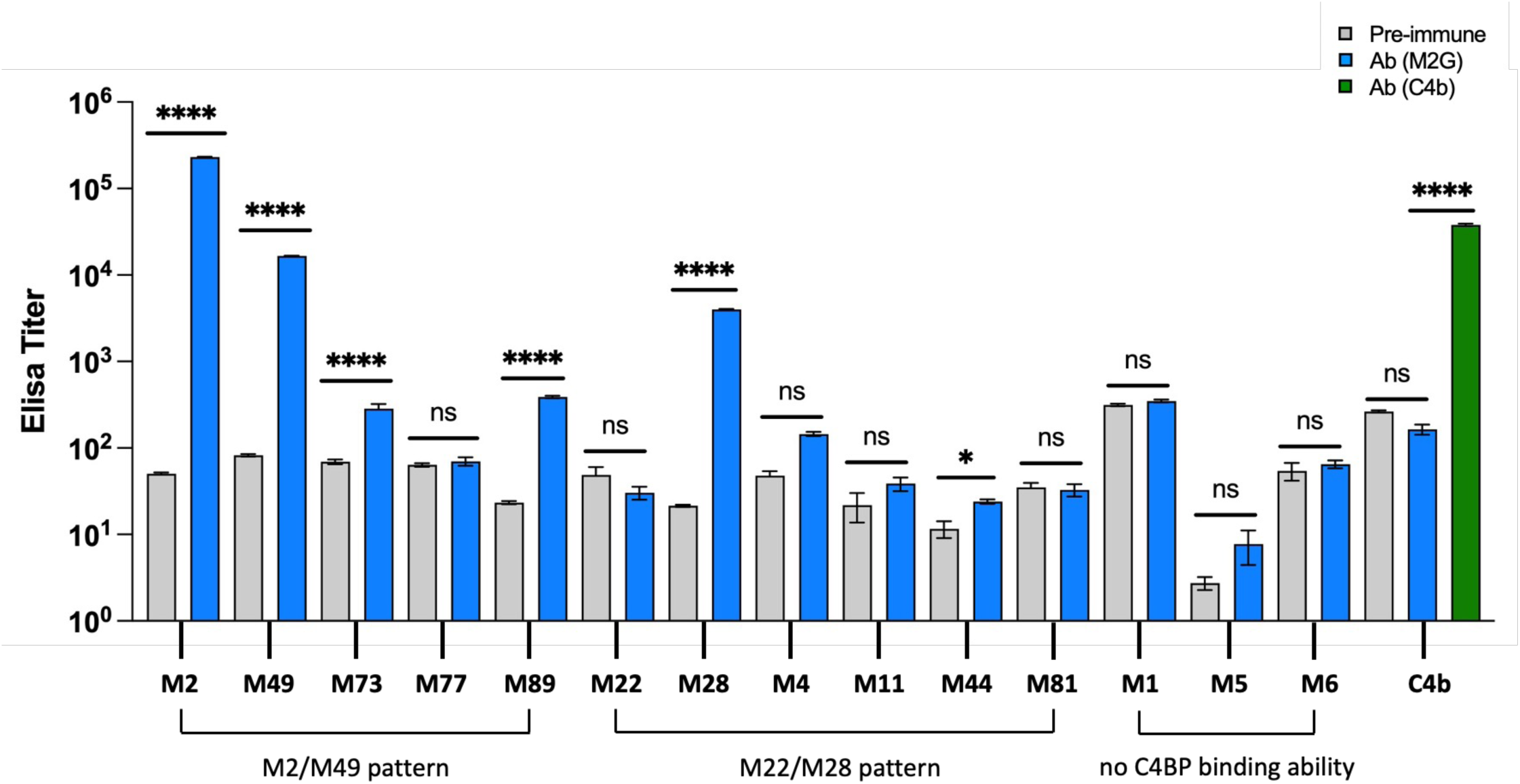
Reactivity and cross-reactivity of M2G antisera from second rabbit. Data are presented as in Figure 3.

**Figure S7.**
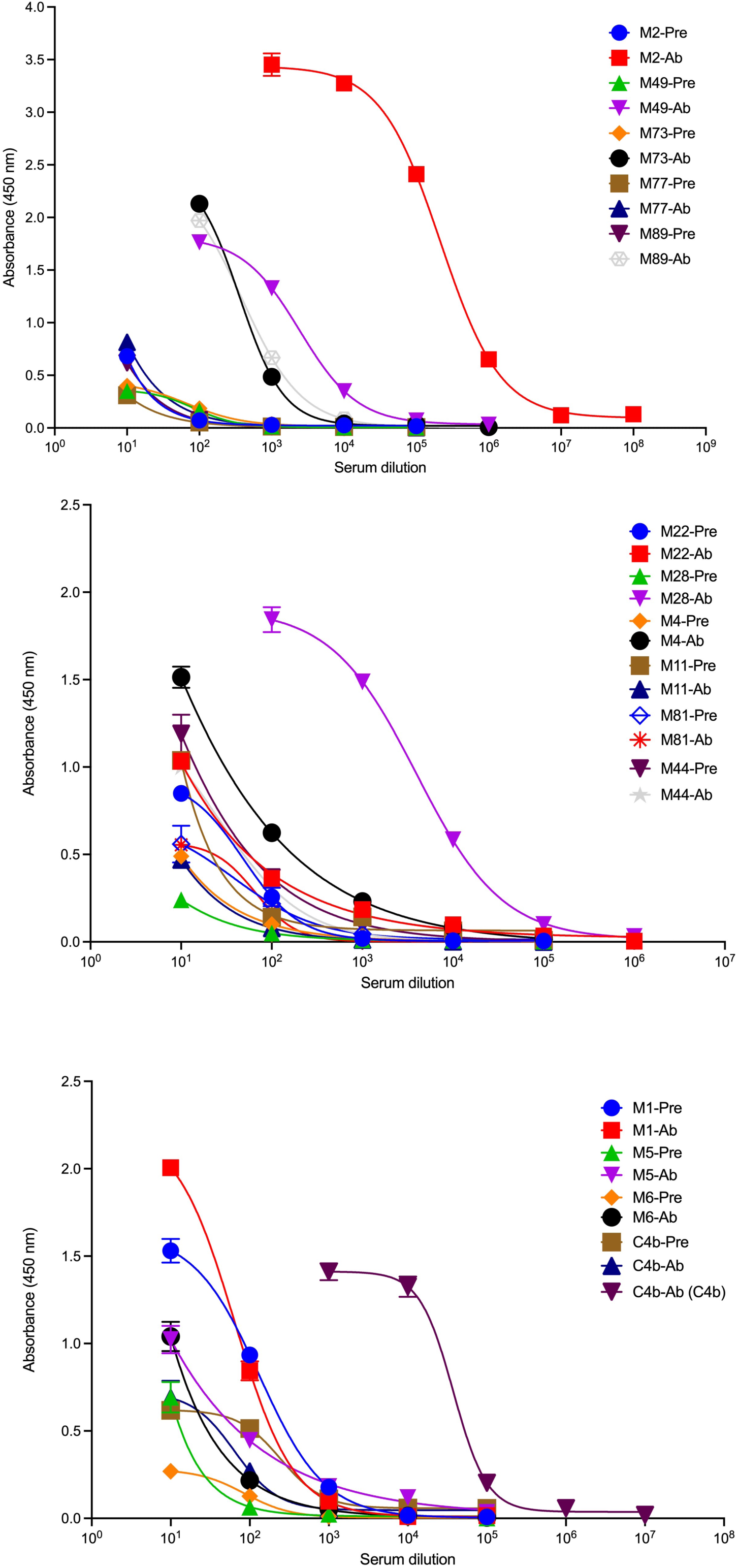
ELISA titers of rabbit sera against M proteins. Data are presented as in Figure S4.

**Figure S8.**
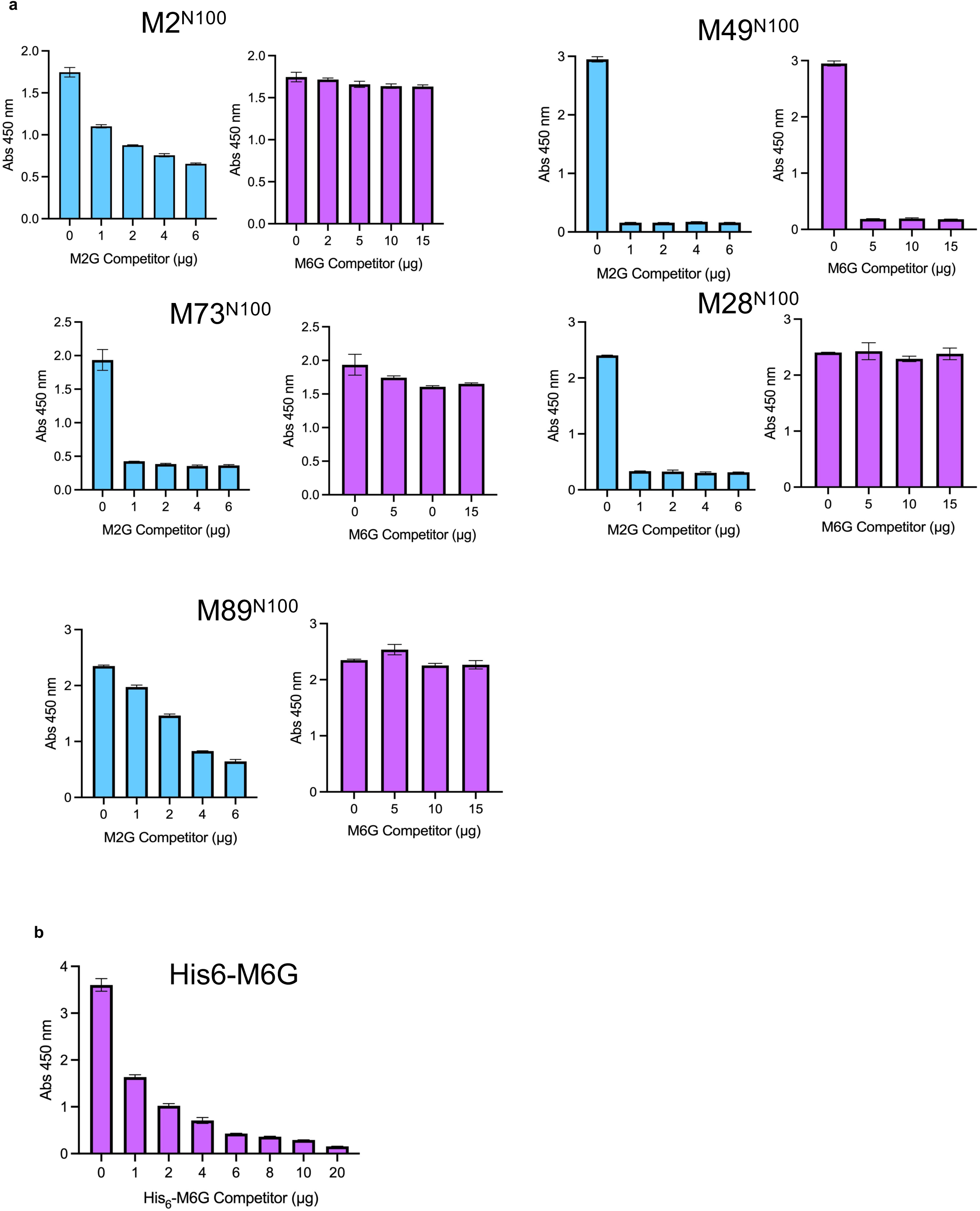
M2G and M6G competition against M2G antiserum. **a.** Binding as determined by ELISA of M2G antiserum to M2^N100^, M49^N100^, M73^N100^, M28^N100^, and M89^N100^, which were adhered to the wells of ELISA plates, was competed with increasing concentrations of M2G or M6G. Experiments were carried out in triplicate, and means and standard deviations are shown. **b.** Binding as determined by ELISA of anti-His antibody to His_6_-M6G, which was adhered to wells of ELISA plates, was competed with increasing concentrations of soluble His_6_ -M6G. Experiments were carried out in triplicate and absorbance was measured at 450 nm. The experiment was carried out in triplicate, and means and standard deviations are shown.

**Table S1.**
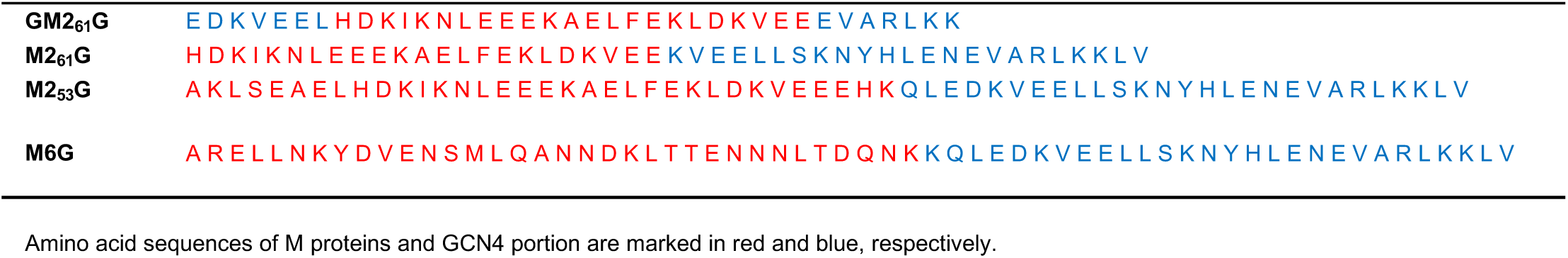
Protein sequences of M Protein-GCN4 fusion constructs.

**Table S2.** Fluorescent intensity of rabbit sera (average of the geometric mean ± standard error)

| **Strep A strains** | **Pre-immune (pre)** | **Ab (M2G)** | **(Ab-pre)/pre** |
| --- | --- | --- | --- |
| M2 | 7.6 ± 0.6 | 205.0 ± 16.0 | 26.1 ± 2.5 |
| M73 | 8.2 ± 1.8 | 28.0 ± 6.0 | 2.4 ± 0.0 |
| M89 | 8.9 ± 0.3 | 13.7 ± 1.0 | 0.6 ± 0.2 |
| M28 | 5.8 ± 1.1 | 14.0 ± 4.4 | 1.3 ± 0.3 |
| M5 | 5.6 ± 0.2 | 6.6 ± 0.4 | 0.2 ± 0.0 |

**Table S3.** Sequence alignment between M2 (aa 53-86) and M proteins (N-terminal 100 amino acids of mature form)

| M2 (53-86) vs M protein | Identity (%) |
| --- | --- |
| M49 | 32.4 |
| M73 | 70.6 |
| M77 | 58.8 |
| M89 | 50.0 |
| M22 | 35.3 |
| M28 | 35.3 |
| M4 | 29.4 |
| M11 | 23.5 |
| M44 | 32.4 |
| M81 | 26.5 |
| M1 | 26.5 |
| M5 | 23.5 |
| M6 | 29.4 |

Sequence alignment was carried out using LALIGN. Cross-reactive M proteins are marked in red.

## Notes

### Competing Interest Statement

The authors have declared no competing interest.

### Summary of Updates

New experiments presented in Figures 5, S6, S7, and S8.

